# Boltzdesign1: Inverting All-Atom Structure Prediction Model for Generalized Biomolecular Binder Design

**DOI:** 10.1101/2025.04.06.647261

**Authors:** Yehlin Cho, Martin Pacesa, Zhidian Zhang, Bruno E. Correia, Sergey Ovchinnikov

## Abstract

Deep learning in structure prediction has revolutionized protein research, enabling large-scale screening, novel hypothesis generation, and accelerated experimental design across biological domains. Recent advances, including RoseTTAFold-AA and AlphaFold3, have extended structure prediction models to work with small molecules, nucleic acids, ions, and covalent modifications. We present BoltzDesign1, which inverts the Boltz-1 model, an open source reproduction of AlphaFold3, to enable the design of protein binders for diverse molecular targets without requiring model finetuning. By utilizing only the Pairformer and Confidence modules, our method significantly reduces computational costs while achieving outstanding in silico success rates and diversity in binder generation. Optimizing directly on the distogram allows us to shape the probability distribution of atomic distances, rather than adjusting a single structure, steering the design toward sequences that yield robust structures with well-defined energy minima. By leveraging a fully atomic model trained on a wide variety of macromolecules, we can generate diverse heterocomplexes with flexible ligand conformations—a capability not currently matched by existing methods. This approach enables the design of novel protein interactions with potential applications in biosensors, enzyme engineering, therapeutic development, and biotechnological innovations.

## 1 Introduction

Designing protein binders that specifically interact with molecular targets to mediate functions such as catalytic activity, molecular recognition, structural support, and protein–biomolecule interactions is a critical goal in both synthetic biology and therapeutic development [1, 2, 3]. With the recent development of deep learning-based computational methods, the success rate of protein design has increased, helping to facilitate designs that can be experimentally validated with minimal screening [4, 5]. Several approaches have emerged for protein design, especially converting structure prediction models into generative models. One strategy is fine-tuning structure prediction models as diffusion models to generate novel structures, exemplified by RfDiffusion[6] and RfDiffusionAA[7]. The alternative strategy employs existing models without additional training by applying backpropagation to iteratively refine sequences until achieving high model confidence scores. This approach, originally termed “hallucination” in the context of trRosetta [8] and later extended to AlphaFold2 [9], has been successfully applied across multiple domains, including fixed-backbone design, binder design, and peptide design [10, 11, 12]. The backpropagation approach offers several significant advantages: it eliminates the requirement for model retraining while permitting optimization based on customized loss functions, such as helical content or radius of gyration. Additionally, it enables modeling of sequence and structure with the unique capability of joint structure prediction of ligand and binder during design, thereby enabling the exploration of induced fit effects upon binding, a feature not currently accessible through computational methods.

BindCraft [10] exemplifies this methodology by utilizing the AlphaFold2 model to generate protein binders through iterative sequence and structure optimization. AlphaFold2 comprises an Evoformer and a structure module, allowing loss to propagate through the entire model to the input. Beginning with relaxed sequence space [13], optimization proceeds toward one-hot encoded sequences, allowing the model to explore solutions that effectively minimize the loss compared to direct one-hot encoded optimization. It leverages output metrics, including confidence scores such as pLDDT and pAE values, and iteratively applies gradients to input sequence until a confident structure is predicted. This approach fully exploits the existing model with various customizable optimization loss functions. For instance, helix loss can modulate the percentage of helices and beta sheets during design to increase structural diversity, while radius of gyration (Rg) loss can regulate the compactness of designed structures.

With the advent of AlphaFold3 (AF3) [14] and RoseTTAFold-All-Atom[7], prediction capabilities now extend beyond protein structures to include small molecules, ions, nucleic acids, and post-translational modifications, such as phosphorylation or glycosylation. Since its release, AlphaFold3 has inspired the development of several comparable models, including Boltz-1 [15] Chai-1 [16], Protenix [17], and HelixFold3 [18]. However, a key distinction between AF2 and AF3-like models is that AF3 models incorporate diffusion modules for structure prediction after processing through MSA and Pairformer modules. Backpropagating through the 200 steps of the diffusion model is computationally expensive, requires substantial memory, is likely to lead to vanishing-gradient issues, and represents only a single sample from the probability distribution of the pair features.

Here, we introduce BoltzDesign1, a framework that enables hallucination through Boltz-1, an all-atom prediction model (Figure 1A), by directly optimizing the probability distribution of the pair features via predicted distrogram. Rather than utilizing the complete model, we employ only the Pairformer and Confidence modules to design proteins applicable to diverse biomolecular applications. As an initial demonstration, we benchmark against small molecules tested in RfDiffusionAA and design binders. BoltzDesign1-generated binders exhibit higher in silico success rates and greater structural diversity compared to RfDiffusionAA. We further show that our approach can be used to design metal binders, nucleic acid binders, and post-translationally modified proteins. Due to the nature of the model simultaneously predicting both binder and ligand, we can uniquely account for flexible ligand modeling—unlike existing methods, where ligands remain fixed. This ability is especially useful when the ligand conformation is unknown or unavailable in the PDB. These advancements will greatly expand our capabilities in designing proteins with more complex folds and functionalities, interacting with a wide range of biologically relevant molecules.

**Figure 1:**
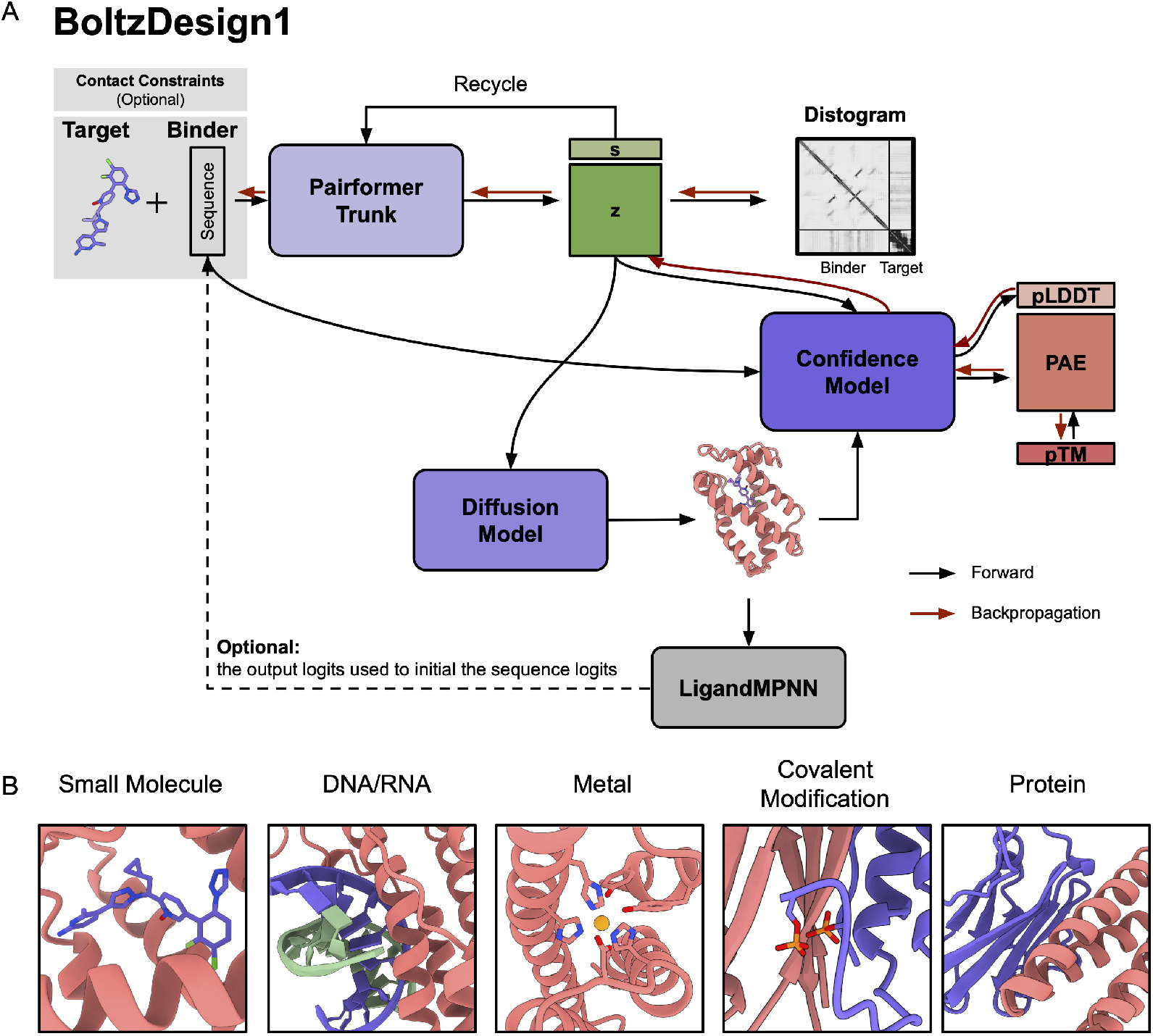
BoltzDesign1 Pipeline. (A) Black arrow is original model prediction pathway and red arrow is direction of loss backpropagation. s represents the single representation, and z represents the pair representation. Loss can be backpropagated through: (1) distogram to Pairformer to sequence, (2) Confidence module to sequence, (3) Confidence module to Pairformer. LigandMPNN is optional. It can either redesign the sequence based on the initial BoltzDesign1-generated structure, or we can initialize the starting amino acid logits from LigandMPNN and further optimize the sequence and structure together. (B) BoltzDesign1 is generalizable to different biomolecules, including small molecules, DNA/RNA, metal, covalent modifications, and proteins.

## 2 Method

The distogram from Pairformer effectively represents the probability distribution of atomic distances that the diffusion model later samples from. By directly optimizing this underlying distribution rather than individual sampled structures, we can achieve more efficient exploration of the sequence-structure landscape while avoiding the computational burden of backpropagating through numerous diffusion steps. Our protein design framework employs two approaches. The simplest method utilizes only the Pairformer to optimize sequences by backpropagating through the distogram loss. A more advanced method incorporates a Confidence module alongside the Pairformer, enabling joint optimization using both distogram loss and confidence scores. We reason that the confidence module represents the fit between the sampled structure from the diffusion module and the probability distribution of the pair features. In some cases, such as the protein-DNA/RNA interactions, where pair features are ambiguous, additional sampling within the diffusion module may better assess the accessible structural space.

To enable backpropagation of the loss from the confidence module to the input, we modify the process to allow gradient flow between the Pairformer and the Confidence module. Specifically, we apply a stop-gradient operation to the Diffusion Module to prevent gradients from flowing through diffusion steps.

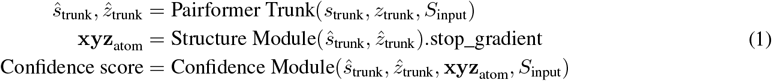

where *s*_*i*_ represents the single-sequence representation, *z*_*ij*_ denotes the pairwise representation, and *S*_input_ is the initial sequence input. **xyz**_atom_ corresponds to the denoised 3D structure coordinates. Here, *ŝ*_trunk_ and 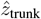 are the outputs from the Pairformer, where gradients are allowed to flow through to the confidence module.

The confidence module receives structure coordinates, updated *ŝ*_trunk_ and 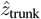 from the Pairformer, but it is initialized with a new representation predicted by the confidence module’s Atom Attention Encoder and MSA module, using the same *S*_input_. In our approach, we backpropagate the confidence loss through the Pairformer to update the sequence representation (S_input).

To optimize a sequence, we implement a four-stage process similar to the optimization approach used in BindCraft.

1. Initially, we explore continuous sequence space using softmax(logits/temperature) with temperature = 1.0 to optimize within distribution constraints, since starting from raw logits often generates unrealistic structures and confidence scores.
2. After the warm-up stage, we initialize with softmax logits and transition to continuous sequence space, where the sequence representation is calculated as (1-*λ*) * logits + *λ* * softmax(logits/temperature), with *λ* = (step+1)/iterations and temperature = 1.0. This allows multiple amino acids to be considered simultaneously at each binder position, enabling broader exploration of the sequence-structure space.
3. In the third stage, we normalize sequence logits to probabilities using the softmax function, gradually converging toward a more realistic sequence representation defined as softmax(logits/temperature). The temperature decreases with each step according to temperature = (1e-2 + (1 - 1e-2) * (1 - (step + 1) / iterations)**2), which also scales the learning rate to decay.
4. In the final stage, we optimize in one-hot encoded sequence space while backpropagating through the softmax representation. The one-hot encoded sequence is given by (argmax(softmax(logits)) - soft-max(logits)).stop_gradient + softmax(logits)
5. Once we obtain the initial BoltzDesign1 generated sequences, LigandMPNN can be utilized for further optimization. This can be done by either fixing the interface or fully redesigning the sequence given a complex structure.

## 3 Results

### Distogram from Pairformer captures protein and target interactions and serves as a proxy for confidence scores

Since backpropagating through the structure (which involves 3D atom coordinates) is computationally expensive, we evaluate whether the distogram is sufficient to achieve good results for both inter- and intra-chain interactions. We first evaluated how much contact information the Pairformer can obtain without using the structure module. We assessed binders from the BindCraft paper by comparing the distogram contacts from Pairformer, the contacts from predicted structures generated by the structure module, and the actual contacts from the AF2-predicted models (Figure 2A). As shown in Figure 2B, Pairformer correctly predicts most of the designs, with 76% of designs having Precision at K(P@K) higher than 0.5. Recycling Pairformer multiple times improves P@K up to three recycling steps, but no significant improvement is observed with additional recycling (Figure 2C).

**Figure 2:**
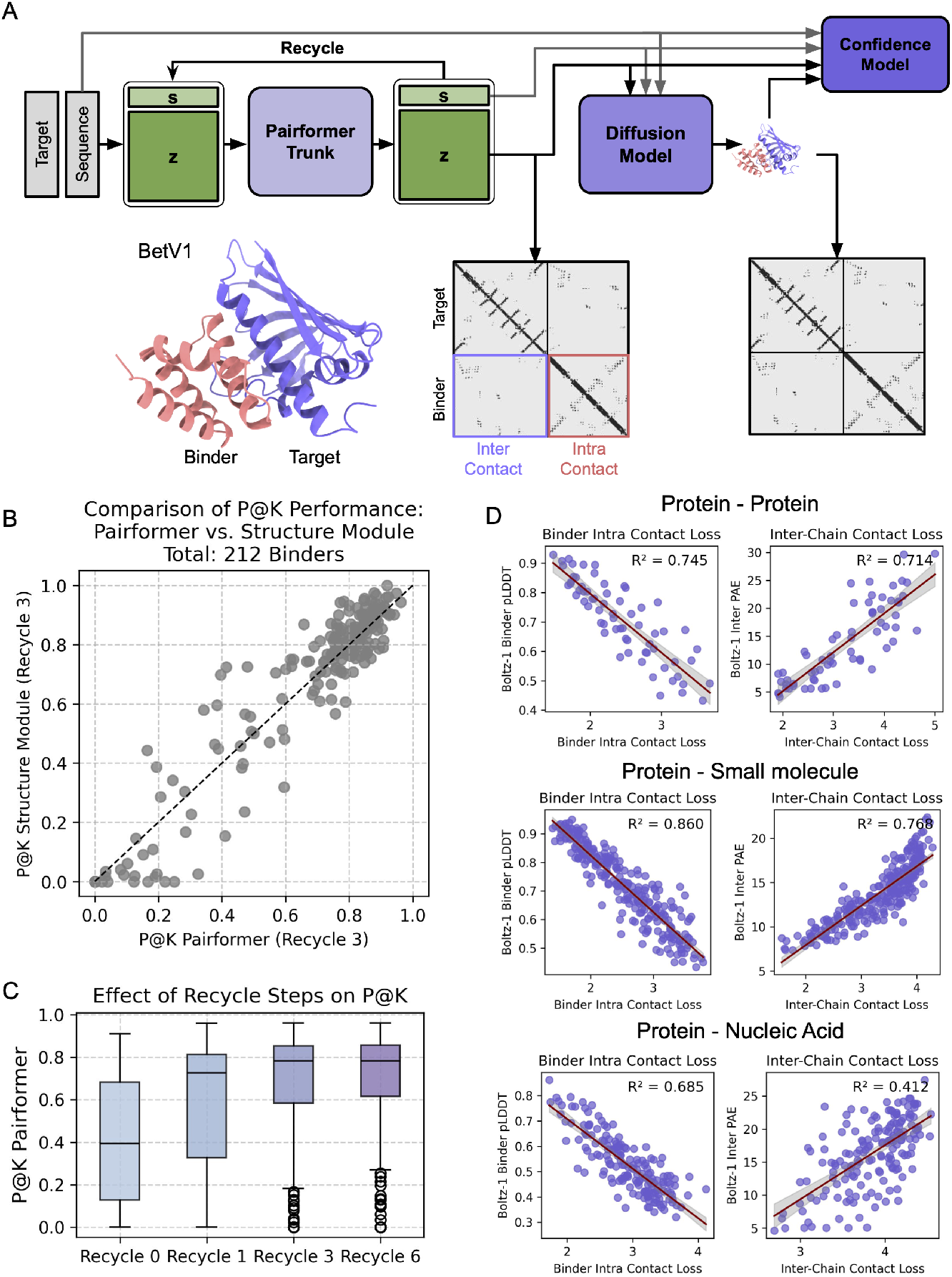
Pairformer distogram effectively captures interactions between proteins and their targets, serving as a proxy for confidence scores. (A) There are two ways of getting contacts. One is from Pairformer, by getting distogram (probability of distance between pairwise residues), and the other is from the diffusion model by obtaining 3D coordinates. Distogram is computed through the Pairformer with the default setting allowing 3 recycles. The sequence and pair representations from Pairformer are passed to the diffusion model, and through 200 diffusion steps, the 3D atomic coordinates structure is achieved. Here, we show the example of BefV1 binder 7 where contacts are correctly predicted by both Pairformer and diffusion model. (B) P@K performance on 212 successful binders across 13 protein targets from the Bindcraft paper. P@K is computed by comparing true inter-chain contacts with predicted inter-chain contacts from Pairformer Distogram and the Structure module’s predicted structures, where K represents the number of true inter-chain contacts (CB-CB distance < 8Å). (C) P@K performance of Pairformer Distogram contacts, comparing the results across different recycling steps (ranging from 0 to 6). (D) Correlation between distogram contact loss and the confidence score of designed binders. From left to right: binder intra-contact loss vs. binder pLDDT, inter-chain contact loss vs. inter-pAE. The rows (from top to bottom) represent protein binders, small molecule binders, and nucleic acid binders. The protein targets are from Bindcraft, while the targets for small molecules and nucleic acids are from the LigandMPNN test dataset.

Next, we examined whether the contact loss from the distogram correlates with confidence scores pLDDT (predicted local distance difference test) and inter-pAE (i-pAE, inter predicted aligned error) to determine if the distogram alone provides sufficient information about structural confidence. We used two contact losses: inter-contact and intra-contact, where inter-contact loss measures the contact between the target and the binder, and intra-contact loss measures contact within the binder. We followed the standard losses from previous hallucination studies [10].

To calculate the contact loss between two residues, we minimize the entropy of low distance bins *p*_*i,j*_ between residues (*i, j*) by calculating

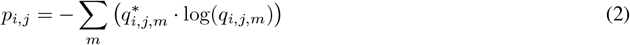

where:

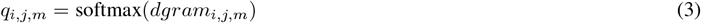

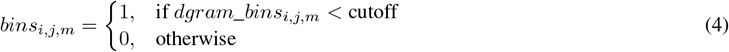

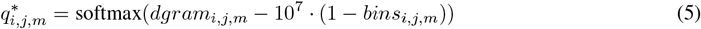

Distogram (*dgram*_*i,j,m*_) encodes the predicted probability of the distance between residues *i* and *j* falling into the *m*-th distance bin. The bins are evenly spaced, ranging from 2 to 22 Å (bin index *m* = 1, …, 64). For each bin *m, dgram*_*bins*_*i,j,m*_ represents the corresponding distance range for that bin. A residue pair is considered a contact if its predicted distance falls below a certain cutoff. For intra-contact, we use a 14 Å cutoff, and for inter-contact, we use a 22 Å cutoff.

To compute overall contact loss, we first compute the mean of the lowest *k* contact losses for each binder residue *i*. For inter-contact loss, we set *k* = 1, meaning only the strongest contact matters. For intra-contact loss, we use *k* = 2, accounting for two key contacts per binder residue. Next, we compute the overall contact loss by averaging the lowest losses from *l* binder residues, where *l* is the length of the binder by default. Additionally, for intra-contact loss, we ignore residue pairs where *i* − *j <* 9. This prevents the model from achieving high confidence simply by generating helical regions, and instead encourages the modeling of long-range interactions.

By analyzing how contact loss relates to pLDDT and inter-pAE, we can determine whether contact-based metrics alone provide enough information to evaluate structural confidence. A strong correlation would suggest that contact loss could serve as an independent measure of protein structure reliability. We designed binders for proteins, small molecules, and nucleic acids, comparing inter- and intra-contact loss with pLDDT and pAE scores. As shown in Figure 2D, we observed a clear correlation between intra- and inter-contact loss and inter-pAE and inter-pLDDT scores for proteins and small molecules. However, for nucleic acids, the correlation between inter-pAE and inter-chain contact loss is less clear, with an R^2^ of 0.412.

For protein-protein interactions and small molecules, distogram contact losses can serve as a proxy for confidence scores. However, for nucleic acids, additional support from confidence losses is required due to the lower correlation between distogram losses and confidence scores. We conclude that while the distogram is sufficient for obtaining good contact information, the structure module is essential for accurately docking biomolecules in more complex and intricate interactions.

### BoltzDesign1 generates diverse small molecule binders with high in silico success rates

RfDiffusionAA paper presented the in silico success rate of binder design for four small molecules: IAI, FAD, SAM, and OQO (Figure 3A). For baseline testing, we also generated binders for these ligands and redesigned them using LigandMPNN [19] to compare our success rate with those reported in RfDiffusionAA. In addition to fully redesigning sequences, we also explored a strategy of fixing binding interface residues within 6Å (CA) of any ligand atom while redesigning surface residues to see whether the initial sequence designed from BoltzDesign1 can model better interfaces than LigandMPNN redesigned interfaces.

**Figure 3:**
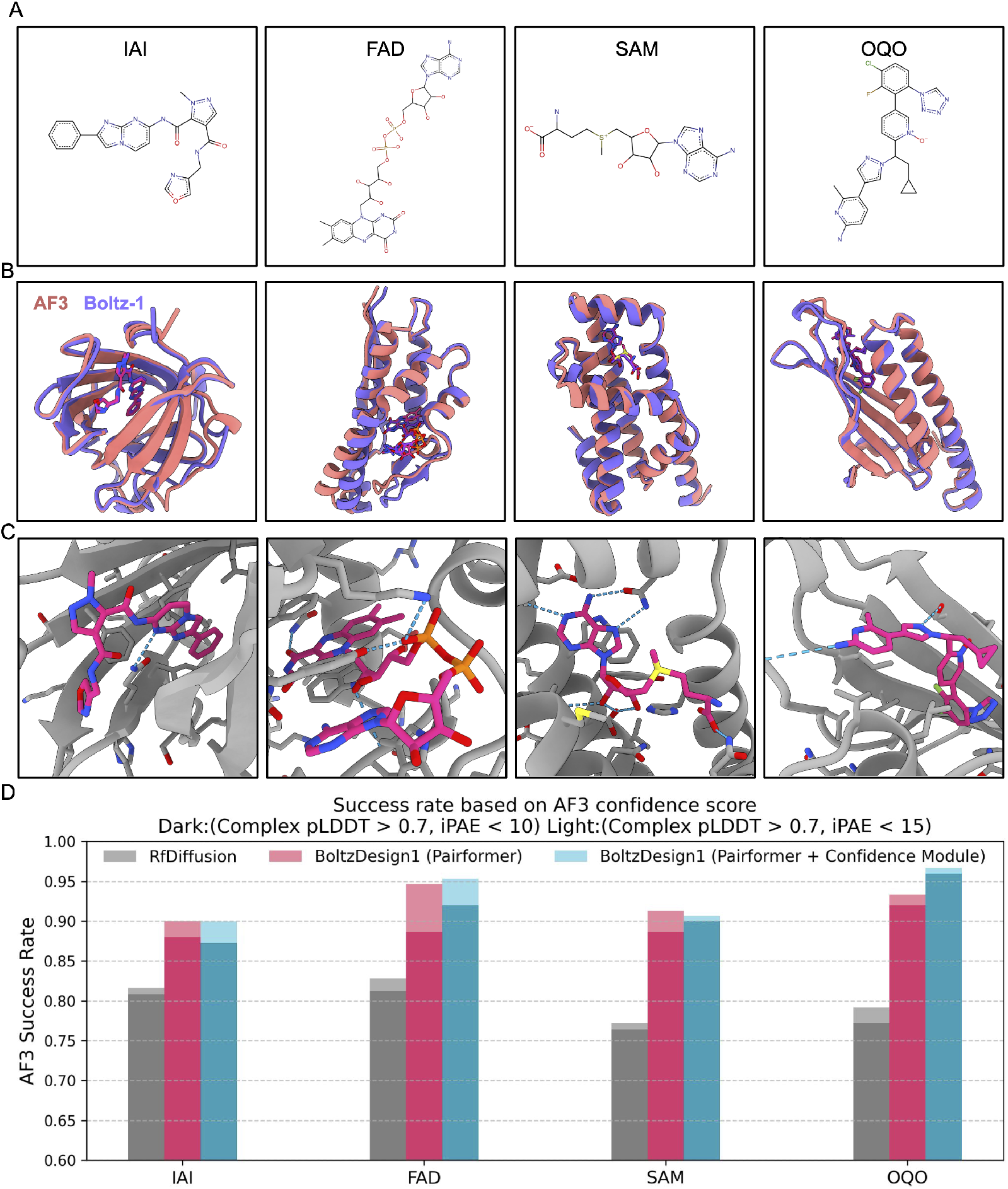
Small molecule binders designed by BoltzDesign1 are highly designable and achieve high AlphaFold3 success rates. (A) Four small molecule candidates validated from the RfDiffusionAA paper: IAI, FAD, SAM, and OQO. (B) Examples of designed binders for IAI, FAD, SAM, and OQO from left to right. (C) Zoomed-in binding pockets, where the blue dashed line indicates a hydrogen bond between the ligand and the protein. (D) Bar plots showing the success rates of designs. Success rates were calculated by measuring the percentage of designs passing either a strict filter (AF3 3complex pLDDT > 0.7 and AF3 ipAE < 10) or a more relaxed filter (AF3 3complex pLDDT > 0.7 and AF3 ipAE < 15). We evaluated BoltzDesign1-generated binders under two different conditions: (1) hallucination with both the Pairformer and the confidence module, and (2) using the Pairformer only. In both conditions, we set recycle = 0, fixed the initial BoltzDesign1 interface residues, and redesigned the surface residues using LigandMPNN.

Inter-contact loss for protein-protein interactions has previously been defined as optimizing for one contact per binder residue (where *k*=1, *l* is the length of the binder, as explained in the Method section). However, we frequently observed that for protein-small molecule interactions, the proteins often wrapped around the ligand, leading to high Apo-Holo RMSD due to the maximization of protein-ligand contacts. To address this issue, we first begin optimization without the ligand, allowing the protein to fold unconditionally. Once the secondary structures are established, we then introduce the ligand and optimize the protein-ligand interactions. Instead of calculating loss from multiple contacts, we optimize a single contact at each step—the one with the lowest contact loss (*k*=1, *l*=1). Although we only optimize one contact directly, making that contact confident often requires additional supporting contacts, which are implicitly encouraged during the process. A key distinction between BoltzDesign1 and RfDiffusionAA is that RfDiffusionAA maintains fixed ligand conformations throughout sampling, while BoltzDesign1 adopts new conformations with each iteration.

For each ligand, we generated 30 structures with helix loss weight terms ranging from 0 to -0.3. We then redesigned the sequences with LigandMPNN, generating 5 sequences per structure, and repredicted complex structures with AF3.

We evaluated the results based on five different criteria:

1. Structure confidence scores: pLDDT for overall complex structure confidence and ipAE for binder and small molecule interface confidence. Success rate is defined by AF3 complex pLDDT > 0.7 and AF3 ipAE < 10 [20, 21, 22].
2. Cross-model consistency: Measured by RMSD between predicted structures from Boltz-1 and AF3 to assess the consistency of designed binders across different models, with a success rate defined by RMSD <2 Å.
3. Self-consistency (Designable backbone): RMSD of protein structures before and after redesign with Boltz-1/AF3 with a success rate defined as RMSD < 2 Å [23].
4. Docking score: Evaluated using Gnina [24, 25], which estimates binding affinity and pose quality for the designed binder and target protein [26]. Docking was performed at the AF3-predicted binding site, and the CNN VS score was calculated by multiplying the CNN affinity and pose scores. Results are compared to the wild-type PDB and ligand to assess binding strength and pose quality.
5. Structure diversity: Measured by the average pairwise TM-score between structures generated for the same ligand target [27].

As shown in Figure 3B, BoltzDesign1 successfully generated binders for four small-molecule targets. These binders formed interactions between proteins and ligands, including hydrogen bonds, hydrophobic interactions, and *π*-*π* interactions (Figure 3C).

Both approaches—(1) Pairformer only and (2) Pairformer with Confidence hallucination—demonstrated higher AF3 success rates compared to RfDiffusionAA designs across all four ligands (Figure 3D). Additionally, using both Pairformer and the Confidence module resulted in a higher success rate than using Pairformer alone for three out of four ligands. We achieved the best results by setting the Pairformer recycling step to 0 and fixing the initial BoltzDesign1 sequence at the interface while redesigning the remaining surface regions using LigandMPNN.(Supplementary Figure S2B). Even with sequences fully redesigned using LigandMPNN, we observed higher sequence recovery at the interface, indicating it is more conserved (Supplementary Figure S1). This highlights that BoltzDesign1 can effectively model interactions between biomolecules, compared to backbone generation models that do not explicitly consider sequence. Furthermore, BoltzDesign1 achieved higher success rates with a recycling step of 0 compared to recycling 1 (Supplementary Figure S2A). This suggests that simpler optimization, like fixing the interface and minimizing recycling, produces more reliable backbone and sequence designs. We reason that during multiple recycling steps, the model may generate adversarial or suboptimal sequences that initially have low confidence, but are misleadingly boosted to higher confidence via multiple iterations [28].

BoltzDesign1 designs show significantly higher cross-model consistency and self-consistency than RFdiffusionAA, both for designs generated using the Pairformer and Confidence module (Supplementary Figures S3A, B) and those generated solely by Pairformer (Supplementary Figures S4A, B). Overall, the BoltzDesign1 structures are highly designable, showing strong agreement between the AF3 and BoltzDesign1 models. Docking scores measured by Gnina indicate that while most designs score lower than the wild-type PDB complex and its ligand, several BoltzDesign1 designs outperform the native ligand binder—specifically, 9.3% of designs for SAM, 7.3% for OQO, and 4.0% for IAI (Supplementary Figure S3C).

In terms of structural diversity, BoltzDesign1 outperforms RfDiffusionAA by generating more diverse designs. We evaluated 30 designs per small molecule type without filtering (Figure 4A). BoltzDesign1 generated structures with an average pairwise TM-score of 0.36, indicating greater diversity, while RfDiffusionAA designs were more similar to each other, with an average TM-score of 0.46. Additionally, our analysis of secondary structure content revealed that increasing the helix loss term for BoltzDesign1 led to higher beta sheet content (Figure 4B), as illustrated in the example structures (Figure 4C).

**Figure 4:**
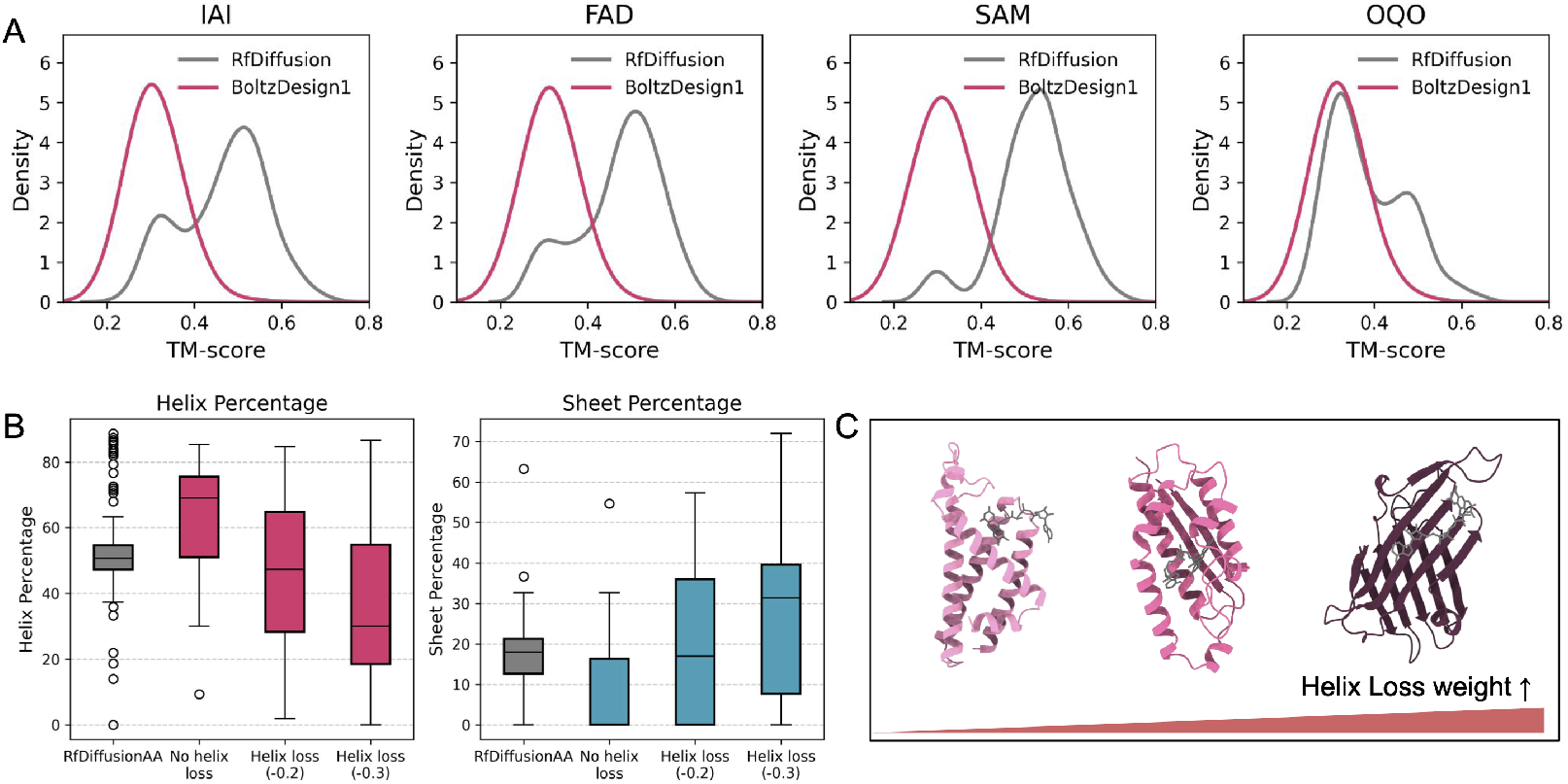
BoltzDesign1 generates highly diverse structures and secondary structure contents. (A) Distribution of pairwise TM-scores for Boltz-1 designs compared to RfDiffusion designs. (B) Secondary structure content of small molecule binders based on different helix loss weights: no helix loss, -0.2, and -0.3 helix loss. (C) Examples of designs showing an increase in sheet content as the absolute helix loss weights increase.

### BoltzDesign1 as a generalizable tool for generating binders to a wide range of biomolecules

Designing binders for biomolecules, including proteins, protein complexes, metal ions, DNA, RNA, and those covalent modifications is crucial for biological pathways and functions [29, 30, 31, 32]. In addition to small molecules, we tested the applicability of BoltzDesign1 to binding other biologically relevant molecules.

Metal ions are essential for enzyme catalysis, signal transduction, and protein regulation, with alkali metals facilitating signal transduction and transition metals like zinc and iron driving enzymatic reactions [33]. As test examples, we selected zinc and iron due to their biological significance and characteristic coordination geometries. We designed binders using the BoltzDesign1 pipeline, redesigned resulting sequences with LigandMPNN, and predicted structures with AF3. These were then analyzed using AllMetal3D to predict metal identity and coordination [34], ensuring that the designs achieved high AF3 confidence scores and a high probability of correct metal identity and coordination. Our designed zinc and iron binders exhibited the expected octahedral and tetrahedral coordination with the protein (Figure 5A,B), incorporating known iron-binding residues such as Tyr, Asp, and His [35].

**Figure 5:**
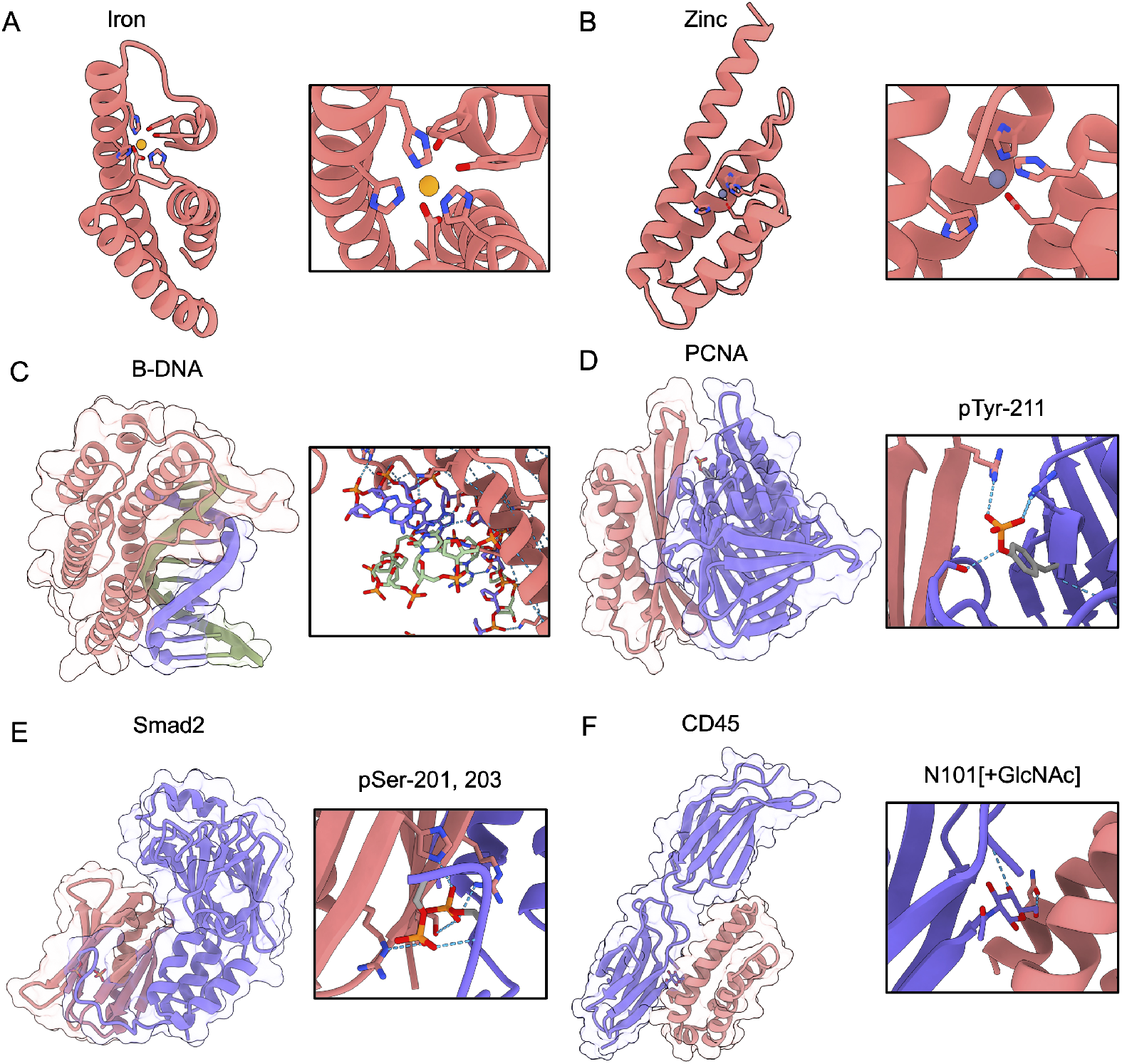
BoltzDesign1 can be used to design sequences and structures that AlphaFold3 predicts to bind to metal ions, nucleic acids, and other biomolecules. (A) Iron and (B) zinc binders. Side chains of designed protein residues interacting with metal ions are visualized. (C) B-DNA binder. purple and green represent the DNA double strand, and pink represents the designed binder. For covalent modifications: (D) PCNA Tyr-211 phosphorylation site binder (E) Smad2 Ser-201, Ser-203 phosphorylation site binder (F) CD45 Asp-101 glycosylation site binder. In panels C-E, pink represents binder proteins, purple represents target proteins, and the blue dotted line indicates hydrogen bond.

Next, for nucleic acid binders, we tested B-DNA [36], the most common natural double-stranded DNA conformation. The binders exhibit high shape complementarity with the nucleic acid, as well as expected positive charge distribution at the interface. Additionally, we observe the formation of relevant hydrogen bonds with both the phosphate backbone and certain base-specific interactions (Figure 5C).

Finally, we demonstrate the design of binders for detecting specific post-translational modifications in target proteins. These covalent modifications play crucial roles in cellular signaling, protein function regulation, and disease progression by altering protein structure, interactions, and localization. We designed binders for therapeutically important proteins that have known phosphorylation or glycosylation sites. The first target is the phosphorylated proliferating cell nuclear antigen (PCNA) at Tyr-211. PCNA-Y211 can promote cancer progression and development [37]. Designing binders that target this site could help mitigate these effects. The second target is phosphorylated Smad2. The C-terminal phosphoserine (pSer) residues of phosphorylated Smad2 induce the formation of a homotrimer, which drives cancer progression. Targeting this phosphorylated site can prevent homotrimer formation [38]. The last target is glycosylated CD45, chosen for its role in regulating T-cell activation, offering the potential to modulate its signaling for anti-tumor therapies. CD45 glycosylation can influence various biological properties, including immunogenicity and specificity, potentially hindering signaling activity [39].

To design binders for these targets, we specified constraints to define binding pocket contacts and covalent modification positions. These constraints were then represented as an additional one-hot encoded token feature provided to the model. As shown in Figures 5 D and E, we successfully designed binders that bind to the PCNA and Smad2 phosphorylation sites by forming hydrogen bonds and also interacting with the target protein. Additionally, binders targeting the glycosylation site of CD45 exhibit specific interactions with the covalently attached sugar moiety (Figure 5).

## 4 Conclusion

In this work, we introduce BoltzDesign1, a computational framework leveraging the Boltz-1 all-atom structure prediction model for protein design without requiring additional fine-tuning. The principal advantage of our approach is the elimination of backpropagation through the computationally expensive diffusion module, significantly reducing resource requirements while maintaining design quality.

Our results demonstrate several significant advances in protein design. First, BoltzDesign1 achieves higher in silico success rates, as measured by AlphaFold3, and generates more diverse structures than RfDiffusionAA. Hallucinating through Pairformer and Confidence module without any recycling steps achieves the best in silico success rate, possibly due to the removal of adversarial and difficult structures that would otherwise require multiple recycling steps for accurate prediction. Second, unlike previous approaches that use fixed ligand conformations, BoltzDesign1 adopts new conformations with each iteration, providing a more flexible framework for complex ligand-protein interactions.

The distogram from Pairformer effectively captures biomolecular interactions and serves as a reliable proxy for confidence scores, allowing us to bypass the computational expense of backpropagating through the structure module while maintaining high design quality. Our strategy of fixing the initial BoltzDesign1-generated sequence at the interface residues while redesigning the surface residues resulted in higher success rates compared to redesigning the entire structure. This underscores the strength of BoltzDesign1 in simultaneously modeling both the structure and sequence at the binding interface. Thus, hotspot conditioning method allows targeted design of binding interfaces, improving specificity and functional performance.

Despite its promising results, BoltzDesign1 has some limitations. It does not currently use templates as input, which could be helpful for guiding the design of structures with high uncertainty or providing constraints for target structures. Additionally, the model lacks integration of nucleic acid MSAs, limiting its ability to effectively design protein-DNA/RNA interactions. Another challenge is the potential for overfitting by using a single model for both design and prediction, which could introduce bias and reduce generalizability, particularly for complex targets or new, unseen structures.

Future work will focus on developing more stable optimization methods tailored to target characteristics and implementing joint optimization between sequence design and structure prediction models. Furthermore, we plan to conduct experimental validation of our computational designs and extend our approach to more challenging targets, including those with high flexibility, such as flexible DNA and RNA, as well as targets with multiple covalent modifications and binders that selectively bind to these modifications. These advances mark a significant step toward generalizable biomolecular design approaches, with potential applications in drug discovery, enzyme engineering, and molecular diagnostics.

## Data and code availability

All data and source code is available at https://github.com/yehlincho/BoltzDesign1.

## Acknowledgements

We appreciate the members of Sergey and Bruno’s labs for their valuable discussions. Y.C. acknowledges financial support from the SBS scholarships and the Takeda Fellowship. S.O. acknowledges funding from NSF grant MCB2032259, and Amgen. We extend our special thanks to Rohith Krishna for providing the RfDiffusion-generated backbones.

## A Method Details

**Figure S1:**
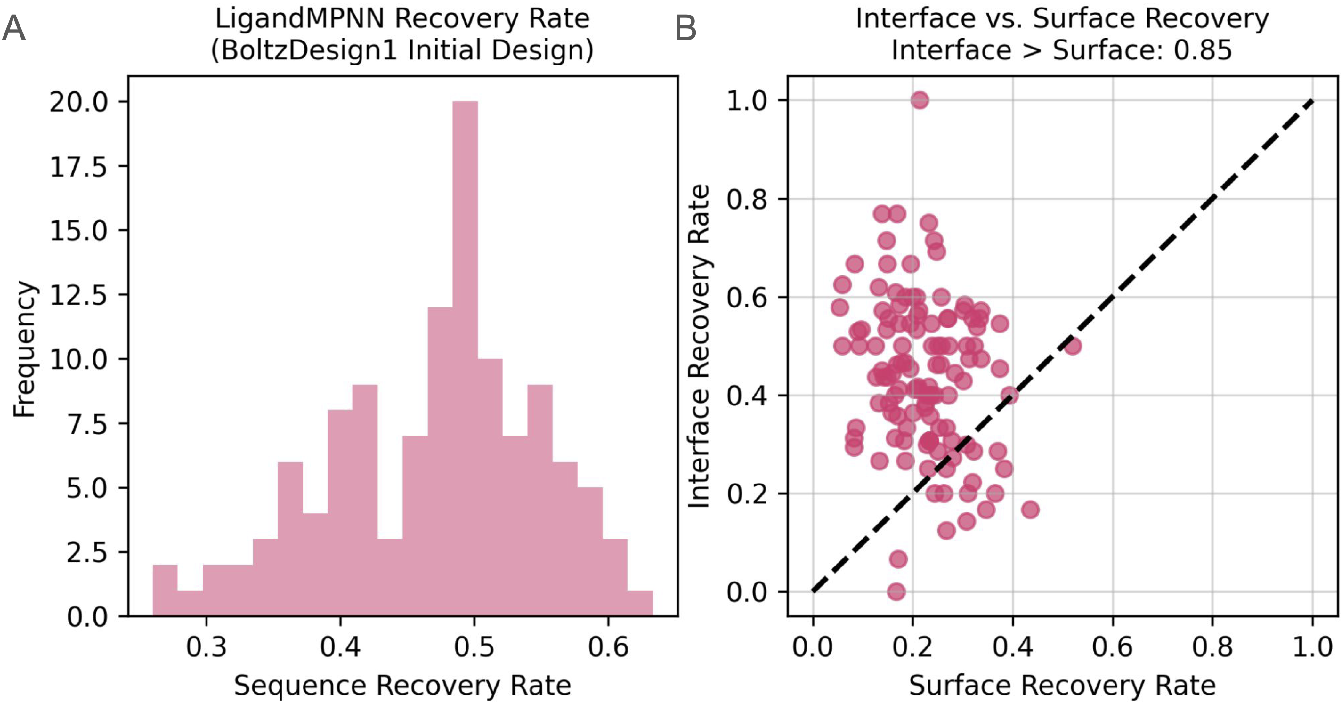
LigandMPNN-designed sequences show a higher sequence recovery rate at the interface compared to surface regions, indicating that the initial BoltzDesign1 interface residues are well-modeled. (A) Overall sequence recovery rate of designs compared to initial BoltzDesign1 sequences. (B) Comparison of sequence recovery rates between surface and interface regions.

**Figure S2:**
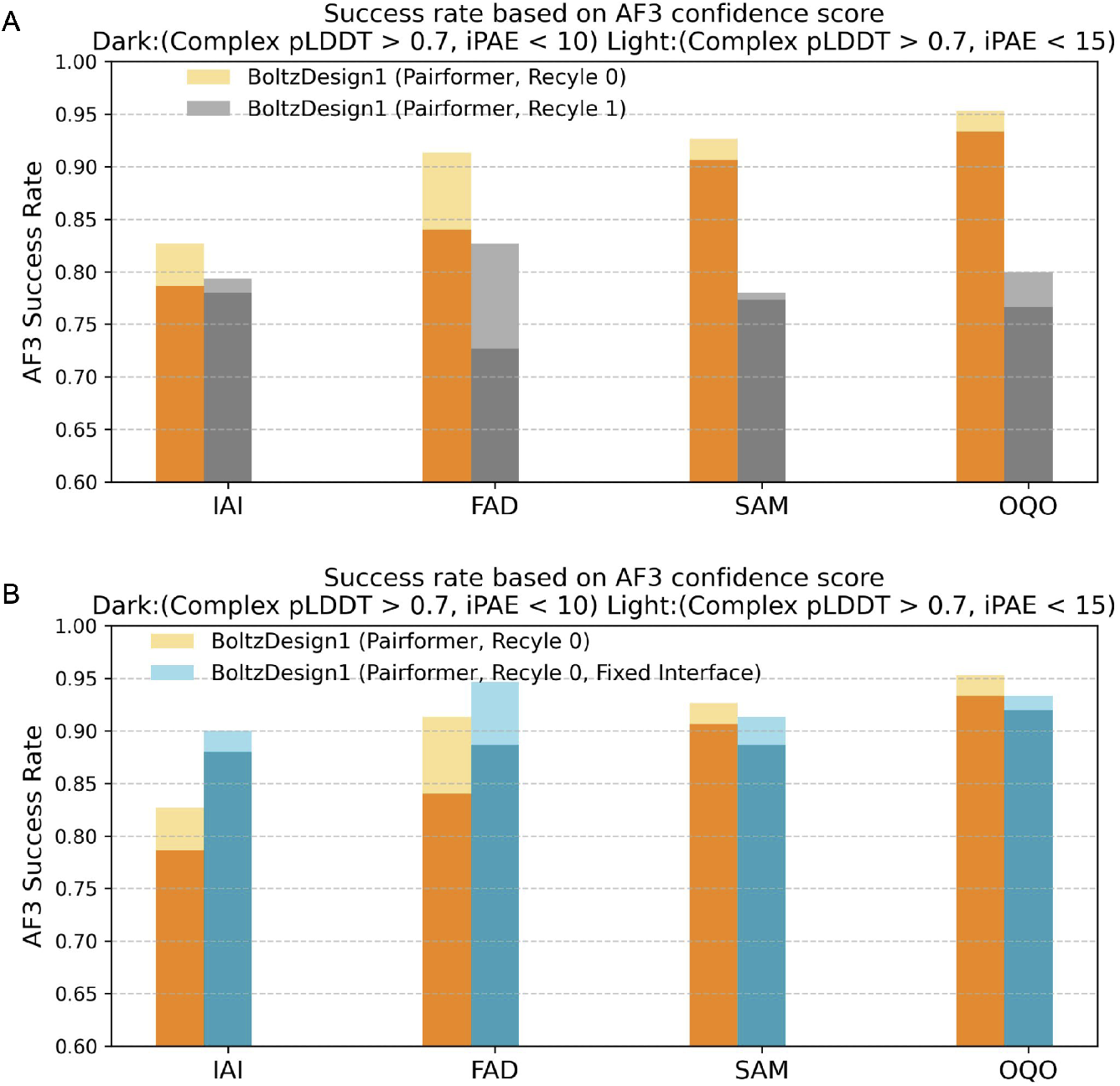
BoltzDesign1 achieves the highest success rate when run without recycling steps, redesigning sequences with LigandMPNN while keeping the initial BoltzDesign1 interface residues fixed. (A) Success rate of BoltzDesign1 (Pairformer) with recycle = 0 vs. recycle = 1. (B) Success rate of BoltzDesign1 (Pairformer) designs fully redesigned with LigandMPNN vs. designs with a fixed interface using the initial BoltzDesign1 sequence and redesigned surface residues using LigandMPNN.

**Figure S3:**
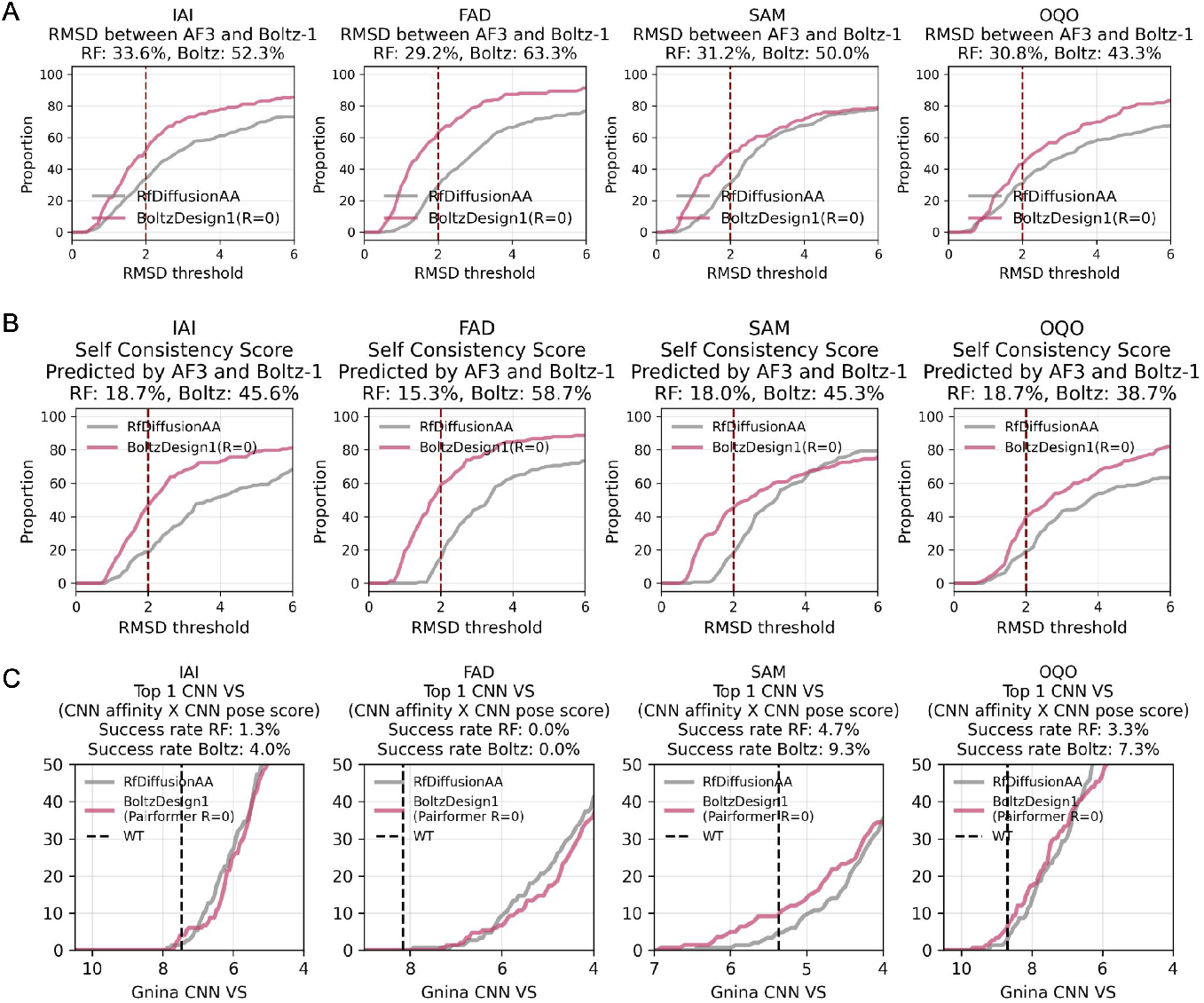
BoltzDesign1 generates binders with highly designable backbones and binding pockets by backpropagating through both the Pairformer and the Confidence module. (A) Agreement between AF3 and Boltz-1 predicted binders measured by RMSD between predictions of the same sequence. (B) Self-consistency score evaluates the quality of backbone structures, showing BoltzDesign1 binders generate highly consistent structures before and after being redesigned with LigandMPNN. (C) Gnina CNN VS scores (CNN affinity × CNN pose score) of the designs, with the dotted line representing the wild-type protein PDB structure and ligand score, where IAI, FAD, SAM, and OQO correspond to the wild-type PDB structures 5SDV, 7BKC, 7C7M, and 7V11, respectively.

**Figure S4:**
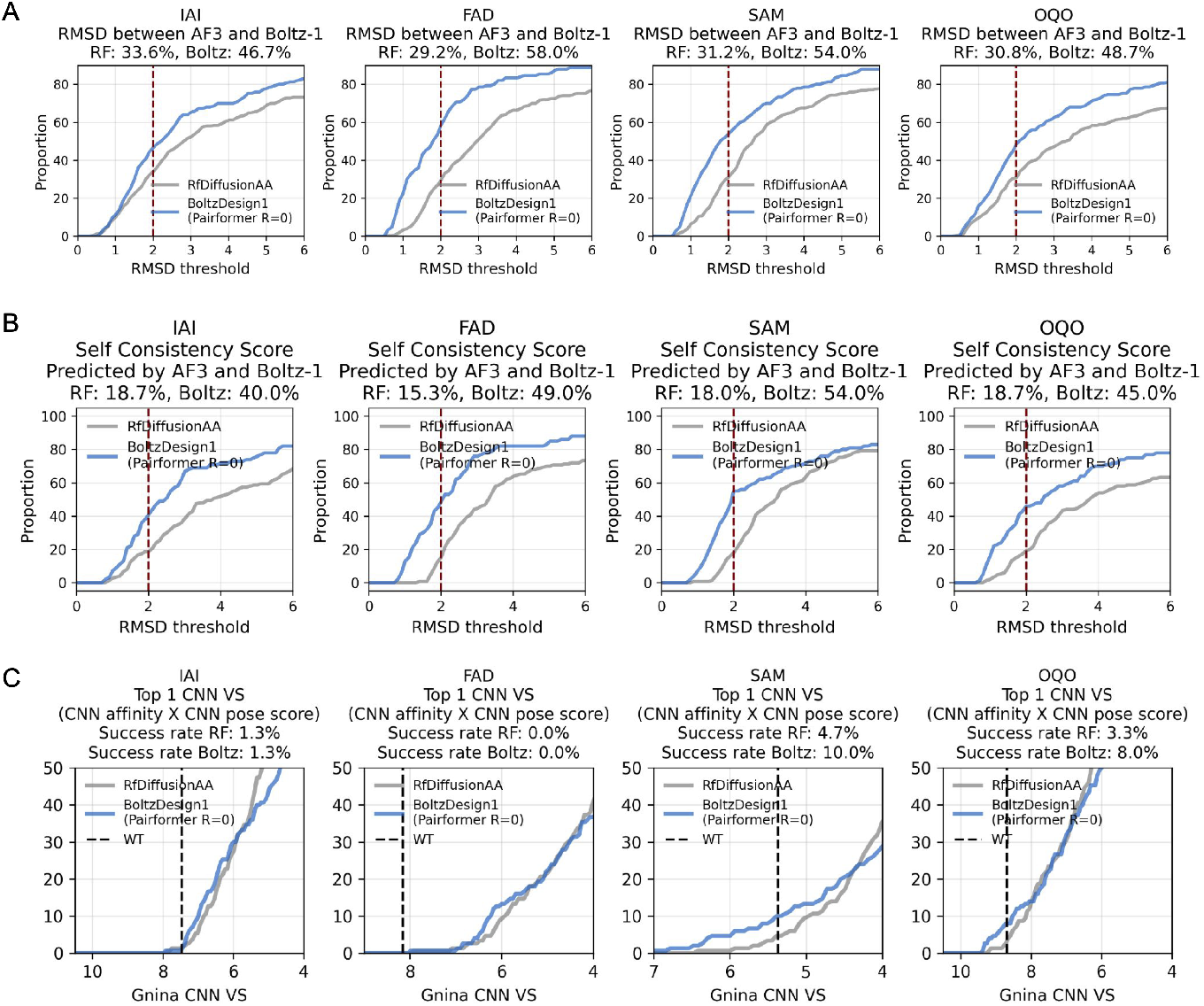
BoltzDesign1 designed binders with Pairformer only and recycle (R=0) in silico scores. (A-C) Evaluated in the same way as described in Figure S3

## References

[1] Zhenyu Yang, Xiaoxi Zeng, Yi Zhao, and Runsheng Chen. Alphafold2 and its applications in the fields of biology and medicine. Signal Transduction and Targeted Therapy, 8(1):115, 2023.

[2] Andrew M Watkins and Paramjit S Arora. Structure-based inhibition of protein–protein interactions. European journal of medicinal chemistry, 94:480–488, 2015.

[3] Haiying Lu, Qiaodan Zhou, Jun He, Zhongliang Jiang, Cheng Peng, Rongsheng Tong, and Jianyou Shi. Recent advances in the development of protein–protein interactions modulators: mechanisms and clinical trials. Signal transduction and targeted therapy, 5(1):213, 2020.

[4] Xingjie Pan and Tanja Kortemme. Recent advances in de novo protein design: Principles, methods, and applications. Journal of Biological Chemistry, 296, 2021.

[5] Hamed Khakzad, Ilia Igashov, Arne Schneuing, Casper Goverde, Michael Bronstein, and Bruno Correia. A new age in protein design empowered by deep learning. Cell Systems, 14(11):925–939, 2023.

[6] Joseph L Watson, David Juergens, Nathaniel R Bennett, Brian L Trippe, Jason Yim, Helen E Eisenach, Woody Ahern, Andrew J Borst, Robert J Ragotte, Lukas F Milles, et al. De novo design of protein structure and function with rfdiffusion. Nature, 620(7976):1089–1100, 2023.

[7] Rohith Krishna, Jue Wang, Woody Ahern, Pascal Sturmfels, Preetham Venkatesh, Indrek Kalvet, Gyu Rie Lee, Felix S Morey-Burrows, Ivan Anishchenko, Ian R Humphreys, et al. Generalized biomolecular modeling and design with rosettafold all-atom. Science, 384(6693):eadl2528, 2024.

[8] Ivan Anishchenko, Samuel J Pellock, Tamuka M Chidyausiku, Theresa A Ramelot, Sergey Ovchinnikov, Jingzhou Hao, Khushboo Bafna, Christoffer Norn, Alex Kang, Asim K Bera, et al. De novo protein design by deep network hallucination. Nature, 600(7889):547–552, 2021.

[9] John Jumper, Richard Evans, Alexander Pritzel, Tim Green, Michael Figurnov, Olaf Ronneberger, Kathryn Tunyasuvunakool, Russ Bates, Augustin Žídek, Anna Potapenko, et al. Highly accurate protein structure prediction with alphafold. nature, 596(7873):583–589, 2021.

[10] Martin Pacesa, Lennart Nickel, Christian Schellhaas, Joseph Schmidt, Ekaterina Pyatova, Lucas Kissling, Patrick Barendse, Jagrity Choudhury, Srajan Kapoor, Ana Alcaraz-Serna, et al. Bindcraft: one-shot design of functional protein binders. bioRxiv, pages 2024–09, 2024.

[11] Casper A Goverde, Benedict Wolf, Hamed Khakzad, Stéphane Rosset, and Bruno E Correia. De novo protein design by inversion of the alphafold structure prediction network. Protein Science, 32(6):e4653, 2023.

[12] Takatsugu Kosugi and Masahito Ohue. Solubility-aware protein binding peptide design using alphafold. Biomedicines, 10(7):1626, 2022.

[13] Christopher Frank, Ali Khoshouei, Lara Fuβ, Dominik Schiwietz, Dominik Putz, Lara Weber, Zhixuan Zhao, Motoyuki Hattori, Shihao Feng, Yosta de Stigter, et al. Scalable protein design using optimization in a relaxed sequence space. Science, 386(6720):439–445, 2024.

[14] Josh Abramson, Jonas Adler, Jack Dunger, Richard Evans, Tim Green, Alexander Pritzel, Olaf Ronneberger, Lindsay Willmore, Andrew J Ballard, Joshua Bambrick, et al. Accurate structure prediction of biomolecular interactions with alphafold 3. Nature, 630(8016):493–500, 2024.

[15] Jeremy Wohlwend, Gabriele Corso, Saro Passaro, Mateo Reveiz, Ken Leidal, Wojtek Swiderski, Tally Portnoi, Itamar Chinn, Jacob Silterra, Tommi Jaakkola, et al. Boltz-1: Democratizing biomolecular interaction modeling. bioRxiv, pages 2024–11, 2024.

[16] Chai Discovery team, Jacques Boitreaud, Jack Dent, Matthew McPartlon, Joshua Meier, Vinicius Reis, Alex Rogozhonikov, and Kevin Wu. Chai-1: Decoding the molecular interactions of life. BioRxiv, pages 2024–10, 2024.

[17] ByteDance AML AI4Science Team, Xinshi Chen, Yuxuan Zhang, Chan Lu, Wenzhi Ma, Jiaqi Guan, Chengyue Gong, Jincai Yang, Hanyu Zhang, Ke Zhang, et al. Protenix-advancing structure prediction through a comprehensive alphafold3 reproduction. bioRxiv, pages 2025–01, 2025.

[18] Lihang Liu, Shanzhuo Zhang, Yang Xue, Xianbin Ye, Kunrui Zhu, Yuxin Li, Yang Liu, Jie Gao, Wenlai Zhao, Hongkun Yu, et al. Technical report of helixfold3 for biomolecular structure prediction. arXiv preprint arxiv:2408.16975, 2024.

[19] Justas Dauparas, Gyu Rie Lee, Robert Pecoraro, Linna An, Ivan Anishchenko, Cameron Glasscock, and David Baker. Atomic context-conditioned protein sequence design using ligandmpnn. Biorxiv, pages 2023–12, 2023.

[20] Nathaniel R Bennett, Brian Coventry, Inna Goreshnik, Buwei Huang, Aza Allen, Dionne Vafeados, Ying Po Peng, Justas Dauparas, Minkyung Baek, Lance Stewart, et al. Improving de novo protein binder design with deep learning. Nature Communications, 14(1):2625, 2023.

[21] Zander Harteveld, Alexandra Van Hall-Beauvais, Irina Morozova, Joshua Southern, Casper Goverde, Sandrine Georgeon, Stéphane Rosset, Michëal Defferrard, Andreas Loukas, Pierre Vandergheynst, et al. Exploring “dark-matter” protein folds using deep learning. Cell systems, 15(10):898–910, 2024.

[22] M Jendrusch, JO Korbel, and S Kashif Sadiq. Alphadesign: A de novo protein design framework based on alphafold.(2021). Back to, page 2.

[23] Jason Yim, Hannes Stärk, Gabriele Corso, Bowen Jing, Regina Barzilay, and Tommi S Jaakkola. Diffusion models in protein structure and docking. Wiley Interdisciplinary Reviews: Computational Molecular Science, 14(2):e1711, 2024.

[24] Andrew T McNutt, Paul Francoeur, Rishal Aggarwal, Tomohide Masuda, Rocco Meli, Matthew Ragoza, Jocelyn Sunseri, and David Ryan Koes. Gnina 1.0: molecular docking with deep learning. Journal of cheminformatics, 13(1):43, 2021.

[25] Ian Dunn, Somayeh Pirhadi, Yao Wang, Smmrithi Ravindran, Carter Concepcion, and David Ryan Koes. Cache challenge# 1: Docking with gnina is all you need. Journal of Chemical Information and Modeling, 64(24):9388– 9396, 2024.

[26] Andrew T McNutt, Yanjing Li, Rocco Meli, Rishal Aggarwal, and David Ryan Koes. Gnina 1.3: the next increment in molecular docking with deep learning. Journal of Cheminformatics, 17(1):28, 2025.

[27] Yang Zhang and Jeffrey Skolnick. Tm-align: a protein structure alignment algorithm based on the tm-score. Nucleic acids research, 33(7):2302–2309, 2005.

[28] Yehlin Cho, Justas Dauparas, Kotaro Tsuboyama, Gabriel Rocklin, and Sergey Ovchinnikov. Implicit modeling of the conformational landscape and sequence allows scoring and generation of stable proteins. bioRxiv, pages 2024–12, 2024.

[29] John A Tainer, Victoria A Roberts, and Elizabeth D Getzoff. Protein metal-binding sites. Current Opinion in Biotechnology, 3(4):378–387, 1992.

[30] Maria Claudia Villegas Kcam, Annette J Tsong, and James Chappell. Rational engineering of a modular bacterial crispr–cas activation platform with expanded target range. Nucleic Acids Research, 49(8):4793–4802, 2021.

[31] Cameron J Glasscock, Robert Pecoraro, Ryan McHugh, Lindsey A Doyle, Wei Chen, Olivier Boivin, Beau Lonnquist, Emily Na, Yuliya Politanska, Hugh K Haddox, et al. Computational design of sequence-specific dna-binding proteins. bioRxiv, 2023.

[32] Dana Pascovici, Jemma X Wu, Matthew J McKay, Chitra Joseph, Zainab Noor, Karthik Kamath, Yunqi Wu, Shoba Ranganathan, Vivek Gupta, and Mehdi Mirzaei. Clinically relevant post-translational modification analyses—maturing workflows and bioinformatics tools. International journal of molecular sciences, 20(1):16, 2018.

[33] Helmut Sigel and Astrid Sigel. The bio-relevant metals of the periodic table of the elements. Zeitschrift für Naturforschung B, 74(6):461–471, 2019.

[34] Simon L Dürr and Ursula Rothlisberger. Allmetal3d: joint prediction of localization, identity and coordination geometry of common metal ions in proteins. bioRxiv, pages 2025–02, 2025.

[35] Sujata Sharma, Mau Sinha, Sanket Kaushik, Punit Kaur, and Tej P Singh. C-lobe of lactoferrin: The whole story of the half-molecule. Biochemistry Research International, 2013(1):271641, 2013.

[36] Richard Wing, Horace Drew, Tsunehiro Takano, Chris Broka, Shoji Tanaka, Keiichi Itakura, and Richard E Dickerson. Crystal structure analysis of a complete turn of b-dna. Nature, 287(5784):755–758, 1980.

[37] Bo Peng, Janice Ortega, Liya Gu, Zhijie Chang, and Guo-Min Li. Phosphorylation of proliferating cell nuclear antigen promotes cancer progression by activating the atm/akt/gsk3β/snail signaling pathway. Journal of Biological Chemistry, 294(17):7037–7045, 2019.

[38] Jia-Wei Wu, Min Hu, Jijie Chai, Joan Seoane, Morgan Huse, Carey Li, Daniel J Rigotti, Saw Kyin, Tom W Muir, Robert Fairman, et al. Crystal structure of a phosphorylated smad2: Recognition of phosphoserine by the mh2 domain and insights on smad function in tgf-β signaling. Molecular cell, 8(6):1277–1289, 2001.

[39] Michelle L Hermiston, Zheng Xu, and Arthur Weiss. Cd45: a critical regulator of signaling thresholds in immune cells. Annual review of immunology, 21(1):107–137, 2003.

